# The Unexpected Functional Diversity of Photoexcited NAD

**DOI:** 10.1101/2023.10.19.563164

**Authors:** Corinna L. Kufner, Mikołaj J. Janicki, Gabriella G. Lozano, Dimitar D. Sasselov

## Abstract

Despite the vital role of nicotinamide adenine dinucleotide (NAD) as a cofactor in all living organisms, the diversity of its functions is poorly understood. Particularly in interaction with ultraviolet (UV) light, a variety of photorelaxation channels can be accessed, which current models lack to explain. In this work, for the first time, we used picosecond UV pump, mid-infrared (mIR) probe spectroscopy and accurate quantum-chemical calculations to elucidate the ultrafast photodynamics of NAD^+^ and NADH to unify contradictory mechanisms from the past decades in the big picture. We found direct evidence for a long-lived (∼900 ps) charge-separated state in NADH, which has been unobserved previously and results in the parallel population of a fluorescent state. The photochemical pathways demonstrated here open up functions of NAD in chemistry and molecular biology, such as an electron donor, as a FRET agent or as a redox pair switch, which have not been considered previously.

**TOC GRAPHICS:** 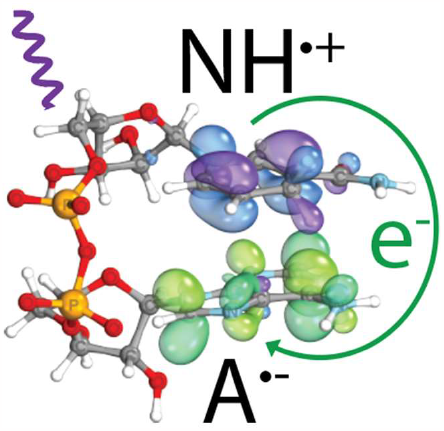

---

Nicotinamide adenine dinucleotide (NAD) is a vital molecule to life: It acts as an electron carrier in all living cells, while the molecule changes its redox state between its oxidized (NAD^+^) and reduced form (NADH).^1-3^ This process is essential to the metabolism of cells and small imbalances in the ratio between the two forms can be indicative of cell disease, such as cancer.^4^ Despite being a vital cofactor, its interactions with ultraviolet (UV) light are not fully understood.^5, 6^ NAD consists of an adenosine (A) and nicotinamide (N) moiety, which are connected by a pyrophosphate backbone (Fig. 1, molecular structures). The addition of a hydride anion (H^-^) at the C4 position of the aromatic pyridine ring (Fig. 1, arrow) alters the photophysical and photochemical properties of the molecule substantially, e.g. it causes a shift of the UV absorbance of the nicotinamide moiety (Fig. 1, red, black) from ∼260 nm to ∼340 nm.^7^ When NAD is a free unbound molecule in an aqueous solution, the chromophores A and N move dynamically between a stacked (folded) and an unstacked (unfolded) conformation.^8-12^ In an aqueous solution at room temperature of 22°C, the fraction of stacked NADH has been reported to be 20 - 55%.^9, 12-16^ In the stacked conformation, the two chromophores can strongly interact with each other through π-stacking forces, which become considerable when the distance between A and N is less than 8 Å.^5, 12, 17^ Surprisingly, the fluorescence lifetimes of NADH have been reported to have longer lifetimes in the stacked than in the unstacked conformation. ^14-16, 18^ The explanations offered for this unusual behavior range from reversible excited-state reactions to complex kinetics, but are not free of contradictions.^6, 13, 19-21^ In 1981, the formation of an exciplex, or a charge-transfer (CT) excited state complex, was proposed,^14^ but there is still no direct proof of this.

**Figure 1.**
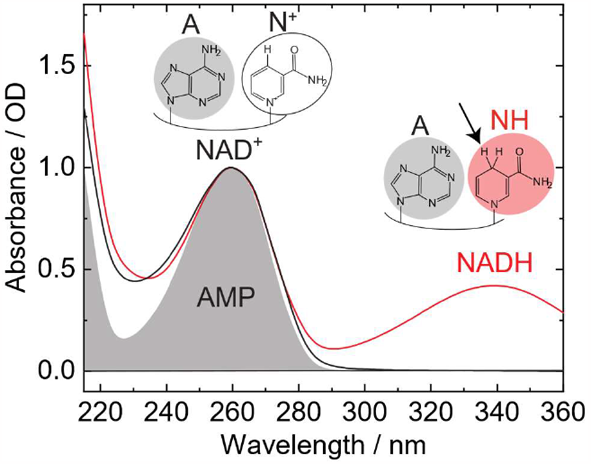
Molecular structures and UV/Vis absorbance spectra of an aqueous NAD^+^(black) and NADH (red) solution at 23°C, 50 mM phosphate buffered (pH 6.9). The absorbance of adenosine monophosphate (AMP) as a function of wavelength is shown as grey shaded area for comparison. All concentrations were adjusted for an absorbance of OD 1.0 at 260 nm for comparison.

Here, we used picosecond UV-pump mid-IR probe spectroscopy with a temporal resolution ranging from several picoseconds up to nanoseconds to determine the ultrafast dynamics after direct photoexcitation of both NADH and NAD^+^. In contrast to previous studies, we probed the vibrational modes for an extended time window, which allowed us to monitor the population of the excited states directly. We also performed accurate quantum-chemical calculations to characterize the photophysical properties and conduct a vibrational frequency analysis of the title molecule. Our findings provide the first direct evidence for the long-lived CT excited state in NADH and none or very short-lived states (< 10 ps) in NAD^+^.

The distinct absorption bands of NADH allowed us to selectively photoexcite the dihydronicotinamide (NH*) moiety at 339 nm and the adenosine (A*) moiety at 267 nm (Fig. 2, left) from their separate electronic ground states. The direct 339 nm irradiation of NADH leads to the population of the bright locally-excited (LE) 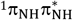 state (Table 1, and Fig. S5 in ESI), resulting subsequently in initiating photochemistry only on the NH fragment. To investigate the photoinduced dynamics of NADH occurring after selective 339 nm excitation, we obtained the transient mid-infrared absorbance difference spectra (Figure 3A, top) of the photoinduced processes in a color map representation as a function of wavenumbers (x) and pump-probe delay times (y). Negative signals are shown in blue and positive signals are in red. The blue bands at 1551 cm^-1^ (antisymmetric two C=C bonds stretching), 1627 cm^-1^ (C=O bond stretching), and 1685 cm^-1^ (symmetric two C=C bonds stretching) are characteristic of the ground state bleach (GSB, Table S3 in ESI) of the dihydronicotinamide moiety (NH).^11^ Global fitting analysis revealed a predominantly mono-exponential decay with a reaction intermediate with a lifetime of 460 ± 150 ps in good agreement with the previously reported slow fluorescent decay of the NH* state (Fig. 2, green arrow).^14, 15, 19, 22^ The corresponding decay-associated difference spectrum (DADS) is shown in the second plot of Fig. 3A. The DADS illustrates a long-lived positive band below 1525 cm^-1^ (purple shaded area) that might be a fingerprint of fluorescence from the NH* state. To clarify the long-lived absorption signal, assuming it comes from the lowest-lying excited state according to the Kasha’s rule^23^, we conducted a harmonic vibrational frequency analysis for the microsolvated dihydronicotinamide moiety, taking the optimized 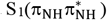 structure found at the ADC(2)/cc-pVTZ level of theory (Fig. S9 in ESI). In the simulated 1510-1750 cm^-1^ IR spectrum in the S_1_ minimum, there is only one major absorption band near 1510 cm^-1^ (Fig. 3A) associated with asymmetric stretching of the amide group. Hence, we have assigned the recorded positive signal below 1525 cm^-1^ to the calculated band near 1510 cm^-1^. Further ADC(2) excited-state potential energy surface exploration of dihydronicotinamide riboside has revealed that the NH* state cannot come back to the electronic ground state in a radiationless and ultrafast manner because of the significant energy barrier of 0.69 eV between the mentioned 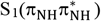 minimum and the corresponding 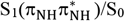 surface crossing (Fig. S7 in ESI). Therefore, the UV-excited dihydronicotinamide moiety, after populating the S_1_ state and due to the inaccessibility of an efficient barrierless non-radiative deactivation pathway, undergoes fluorescence in agreement with previous work.^5, 13, 14, 16, 18, 24, 25^ Furthermore, our multireference MS-CASPT2 calculations predicted that emission from the S_1_ state of the NH^*^ moiety should occur at a wavelength of 475 nm (Fig. S7 in ESI), and it matches very well the recorded maximum fluorescence of around 466 nm in aqueous solution.^14, 15^ Consequently, the discussed experimental-theoretical findings indicate that the recorded absorption signal below 1525 cm^-1^ can be a signature of the long-lived 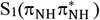 excited state of NADH experiencing fluorescence (Fig. 3A, bottom).

**Table 1.**
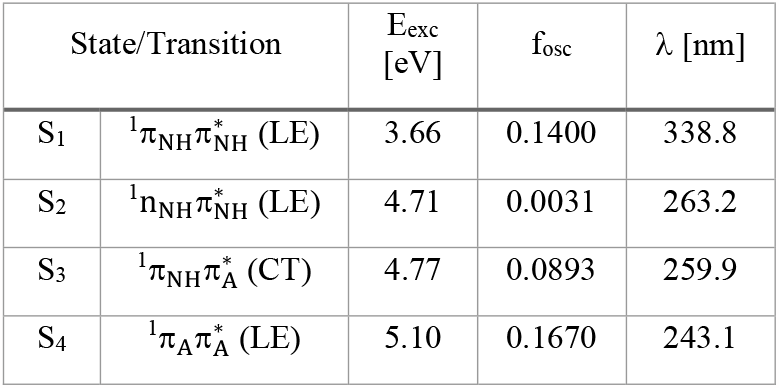
Vertical excitation energies (in eV) of NADH were obtained using the COSMO-ADC(2)/def2-TZVP method, assuming the extracted NADH structure from the solvated system found at the ONIOM(ωB97X-D3/def2-TZVP:GFN2-xTB) level of theory.

**Figure 2.**
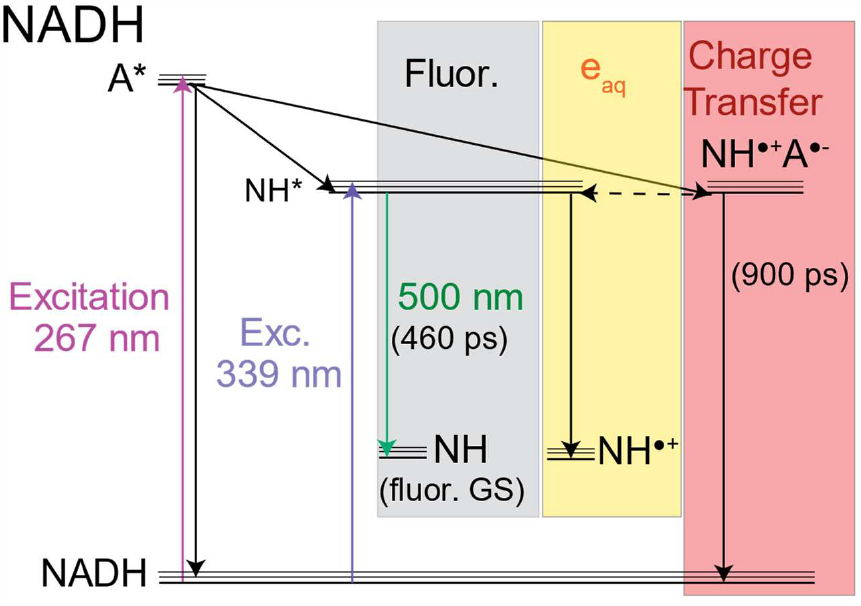
Proposed mechanism for the excited state decays of NADH.

**Figure 3.**
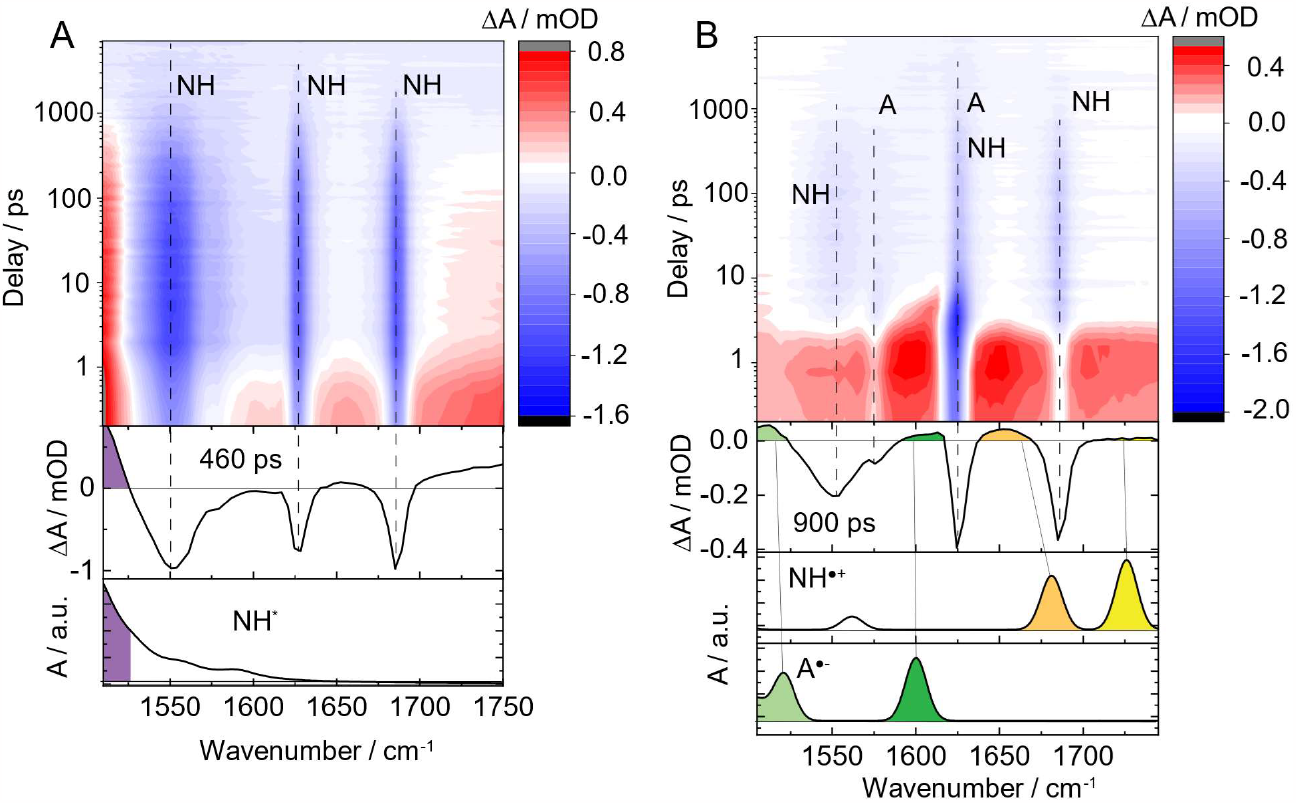
Transient mid-infrared absorbance difference spectra of NADH following (A) 339 nm and (B) 267 nm excitation in aqueous buffered (pD 6.9) solution, as a function of wavenumber (x-axis) and delay time (y-axis). Positive signals are shown in red and negative in blue. The second plot shows the decay associated difference spectra from a global fit analysis, which are associated with several 100 ps lifetimes. The simulated mid-IR spectra for transient reaction intermediates are shown at the bottom: (A) NH* indicative of fluorescence, (B) NH^•+^ and A^•-^ indicative of charge transfer.

After selective excitation of the adenosine moiety (A*) at 267 nm, the direct population of A* occurs through the bright locally excited 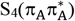 state (Table 1, and Fig. S5 in ESI). In turn, the A* state can be readily deactivated in a radiationless and ultrafast manner from the 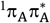 excited state through well-established ring-puckering deactivation pathways of adenosine.^26-28^ Importantly, the initially populated 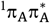 excited state can also serve as a precursor for the consecutive population of the lowest-lying 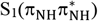 state (Table 1). In the folded form of NADH (Fig. 4A), the latter state can be easily accessible during the rapid non-adiabatic electronic relaxation from the 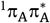 state, according to Kasha’s rule.^23^ Considering previous studies, that discussed mechanism was classified as an ultrafast (<100 fs) intramolecular energy transfer from the adenosine to dihydronicotinamide moiety,^5, 14^ enabling the population of the bright 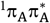 state. Accordingly, apart from the internal relaxation of adenosine, fluorescence from the NH^*^ state should be another deactivation channel of NADH upon 267 nm excitation, and it was experimentally confirmed. ^5, 13, 14, 19, 22^ Surprisingly, our photophysical calculations of the folded NADH molecule (Fig. 4A) predicted the charge-transfer 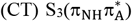 excited state (Table 1, 4.77 eV), in which a virtually single electron (0.74 e^-^) is transferred from the dihydronicotinamide to adenosine moiety. As a result, the CT state could entail a charge separation in NADH, allowing for the formation of the adenosine radical anion and dihydronicotinamide radical cation. Furthermore, the charge-transfer state lies noticeably lower than the bright 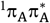 state of adenosine (ΔE = 0.33 eV) (Table 1). Also, it has a non-negligible oscillator strength of 0.089, suggesting that after 267 nm excitation the 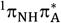 state might be even populated directly or indirectly from the bright 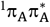 state. Importantly, the inter-ring distance of dihydronicotinamide and adenine moieties and their mutual arrangement in NADH can easily affect the attainability of the CT state.^5^ Therefore, only in the folded form of NADH should the charge-separated state appear. Moreover, the discussed CT state is unexpectedly in the low-energy spectrum (<5.0 eV), which is dominated by locally excited states in short RNA oligonucleotides.^29, 30^ This unique photophysical property of NADH might come from the fact that the NH fragment does not possess an aromatic structure, which energetically stabilizes π-electrons. Thereby, dihydronicotinamide should easier donate an electron than heteroaromatic canonical nucleobases to a nearby chemical environment upon UV excitation.

**Figure 4.**
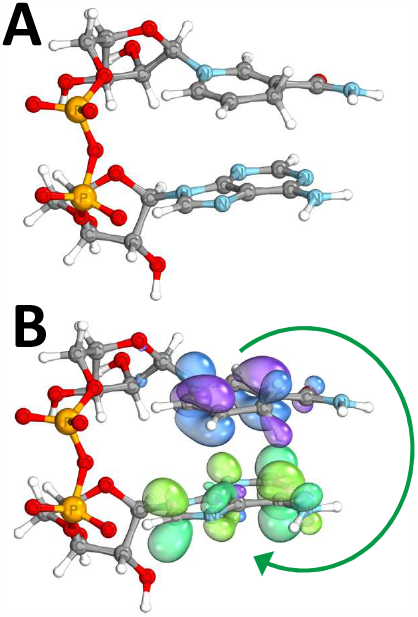
(A) The equilibrium NADH ground-state structure was extracted from the solvated NADH system found at the ONIOM(ωB97X-D3/def2-TZVP:GN2-xTB) level of theory. (B) The presented occupied π_N_ (dihydronicotinamide, purple and blue) and unoccupied 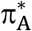 (adenosine, green) molecular orbitals correspond to the charge-transfer 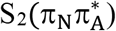 excited state. The green arrow shows a charge transfer of 0.74 e^-^ from the dihydronicotinamide to adenosine moiety.

The experimental absorbance difference spectra of NADH following selective 267 nm excitation of the adenosine moiety (A*) are shown in a color map representation in Fig. 3B. After photoexcitation, characteristic ground state bleach bands of NH (1553 cm^-1^, 1685 cm^-1^: antisymmetric and symmetric two C=C bonds stretching, respectively; 1624 cm^-1^: C=O bond stretching) and A (1575 cm^-1^: C-N/C-NH_2_ bonds stretching, 1624 cm^-1^: ring/NH_2_ stretching) emerge (Table S3 in ESI).^11, 31^ A bi-exponential fit of the individual A and NH GSB bands reveals long (τ_2_) intermediate lifetimes on the order of several 100 ps (1553 cm^-1^(NH): 880 ps, 1575 cm^−1^(A): 280 ps, 1624 cm^-1^ (A, NH): 710 ps, 1685 cm^-1^ (NH): 470 ps). Due to the several coinciding photorelaxation processes, a global fitting analysis is challenging. A 4-exponential fitting approach reveals lifetimes of ∼0.2 ps, ∼10 ps, ∼100 ps and ∼900 ps. The ≤ 10 ps lifetimes can be assigned to coherent artifacts and vibrational cooling in which the excess excitation energy is dispersed to the solvent.^32^ Vibrational cooling limits the time resolution of transient mid-IR spectroscopy, so that the rapid (< 10 ps) energy transfer between the initially excited states cannot be directly observed.^13, 15, 22, 24, 25^ The ∼100 ps and ∼900 ps lifetimes agree with previously published bi-exponential decay times within their error bars of several ±100 ps.^5, 14^-^16, 19, 22, 33^ The DADS corresponding to the ∼900 ps is displayed as the second plot in Fig. 3B. The DADS shows positive bands around 1510 cm^-1^ (light green), 1610 cm^-1^ (dark green), 1654 cm^-1^ (orange), and 1726 cm^-1^ (yellow). To elucidate the origin of these long-lived positive bands, we have assumed that due to a plausible population of the 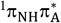 excited state (Table 1), the consecutive charge separation process (Fig. 4B, green arrow) might lead to the formation of the adenosine radical anion (A^•-^) and dihydronicotinamide radical cation (NH^•+^). For this reason, we conducted a vibrational frequency analysis (Table S3 in ESI) using the ωB97X-D3/def2-TZVPPD/CPCM(Water) method for the NH^•+^ radical cation (1562 cm^-1^, 1681 cm^-1^, 1726 cm^-1^) and the A^•-^ radical anion (1520 cm^−1^, 1600 cm^-1^). The resulting simulated mid-IR spectra are shown at the bottom of Fig. 3B, and corresponding experimental and theoretical bands were assigned based on the color coding. In the 1510-1750 cm^-1^ range, our theoretical calculations predicted IR signatures of the NH^•+^ and A^•-^ radicals, which agree well with the recorded long-lived positive signals. Thereby, the obtained results indicate the formation of transient charged radical intermediates that can originate from the population of the charge-transfer state, which allows for intramolecular excited-state electron transfer from the dihydronicotinamide to adenosine moiety upon 267 nm excitation of NADH.

Despite the predominant fluorescent decay channel after 339 nm excitation of NADH, indications of a different photorelaxation pathway can be found in the positive bands around 1653 cm^-1^ and 1725 cm^-1^ of the DADS in Fig. 3A. As described in the previous paragraph, we assigned both long-lived positive bands to the signature of the NH^•+^ radical associated with the charge-separated state. However, in this case, there are no accompanying long-lived fingerprints of A^•-^ radical, which were found upon 267 nm excitation. Hence, these results suggest another active photochemical process occurring after selective excitation of the NH fragment. For the dihydronicotinamide riboside molecule, we performed photophysical calculations (Table 1 and Fig. S6 in ESI) at the ADC(2)/aug-cc-pVDZ level of theory. In the low-energy spectrum, we surprisingly found a repulsive low-lying ^1^πσ^∗^ excited state at 4.27 eV, in which the σ^∗^ orbital is delocalized around both hydrogen atoms at the C4 atom (Fig. S6 in ESI). For this reason, we suspect that the population of the ^1^πσ^∗^ state could enable the photochemical formation of solvated electrons,^34^ which leads to the NH^•+^ radical after 339 nm excitation. Previous time-resolved transient absorption studies of NADH,^35^ showing a signature of solvated electrons after 355 nm excitation appear to confirm this hypothesis. We suppose that upon selective 339 nm excitation, the fluorescent lowest-lying excited state might compete with the ^1^πσ^∗^-mediated solvated electron production during the excited-state dynamics, and the latter state could also contribute to the photochemical events of UV-excited NADH.

Photoexcitation of NAD^+^ at 267 nm yielded the transient mid-infrared absorbance difference spectra in Figure S1 in ESI. Upon excitation, characteristic ground state bleach signatures of A (1573 cm^−1^: C-N/C-NH_2_ bonds stretching, 1622 cm^-1^: ring/NH_2_ stretching) and N^+^ (1662 cm^-1^: C=O bond stretching) emerge in very good agreement with the literature (Table S3 in ESI).^11, 31^ A broad positive (red) band underneath the adenosine GSB ∼1575 cm^-1^ can be observed. All bands decay rapidly, within 7 ± 5 ps, which is indicative of vibrational cooling. We could not observe any signatures of excited states with longer lifetimes in aqueous NAD^+^. The explanation for the difference in populating long-lived excited states between NADH and NAD^+^ might arise from the fact that a small amount (<15%) of NAD^+^ molecules are in the folded form or nicotinamide could have poorer electron-donating properties than its reduced form.^10, 36^ Hence, we hypothesize that any inter-ring excited-state electron transfer in NAD^+^ may have very low quantum efficiency. To gain insight into the population of excited states of NADH and NAD^+^, we conducted an analysis of the GSB signal at 1623 cm^-1^. For comparison, both plots were normalized to the absorbance difference signal at 3 ps (Fig. S2 in ESI). The time-dependence mid-IR data have shown that around 34% of NADH molecules are in excited states over 100 ps, followed by slow decay lasting up to 1000 ps. In the case of NAD^+^, the complete depopulation of excited states occurs within 10-20 ps.

The bi-exponential photochemical relaxation of UV-excited NADH has been the subject of intense studies over the past decades, but its causes remained unclear.^5, 13, 14, 16, 33, 37^ In this work, for the first time, we used transient UV-pump, mid-IR probe spectroscopy and accurate quantum-chemical calculations to solve a well-known puzzle about considerably longer fluorescence reported for aqueous NADH in the stacked arrangement. ^5, 13^-^16, 18, 19, 22, 24, 25, 37^ Our joint experimental-theoretical studies have demonstrated that the elongated fluorescence lifetime can be associated with previously unseen charge-transfer (CT) 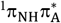 excited state, in which an electron is transferred from the dihydronicotinamide to adenine fragment. Upon selective 267 nm excitation of the adenosine fragment, both long-lived (CT) 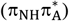 and fluorescent 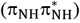 excited states can be initially populated through a non-adiabatic radiationless deactivation mechanism. Then, the former state enables a charge-separation mechanism in NADH, resulting in the formation of long-lived NH^•+^ and A^•-^ intermediates, which we observed in our transient absorption measurements. Subsequently, the part of the collected population in the charge-separated state can overflow to the fluorescent state during the excited-state dynamics, extending the emission lifetime about the time spent in the charge-transfer state which explains the anomalously long fluorescence lifetime. Additionally, the repulsive 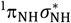 state can mediate the formation of solvated electrons reported by Boldridge et al.^35^ upon selective 339 nm excitation. Interestingly, in the oxidized form, NAD^+^, we have not observed any UV-induced long-lived photochemical processes.

Apart from the mechanistic and spectroscopic results, we demonstrate that dihydronicotinamide riboside has excellent UV-activated electron-donating properties, and it seems to be rare among heterocyclic biomolecules occurring in extant biology, for instance, in RNA oligonucleotides.^29, 30, 38^ UV-excited dihydronicotinamide can also produce solvated electrons,^35^ which can be employed to perform the chemical reduction. Due to the great UV-activated electron-donating properties of the dihydronicotinamide moiety, a single electron can be efficiently and selectively transferred to a nearby acceptor molecule. These properties can be beneficial in a broad range of chemistry. For instance, finding a plausible prebiotic chemical synthesis pathway of dihydronicotinamide riboside could benefit different aspects of the origins of life studies. The NADH/NAD^+^ system can be considered a redox on/off switch as the excitation energy storage at picosecond timescale in the form of the charge-separated state, which can be further used to trigger secondary photochemistry in all living organisms exposed to sunlight. More generally, the long-lived fluorescent state can be used in Förster resonance energy transfer,^15, 39^ which is currently employed in biological and biophysical fields to explain various biochemical processes at the molecular level. As such, dihydronicotinamide riboside appears to be a multifunctional chromophore that naturally occurs as a part of the NAD cofactor in living organisms and could find applications not only in prebiotic chemistry or biological studies but much more. Table 2 shows a summary of the diverse molecular functions of NAD.

**Table 2.**
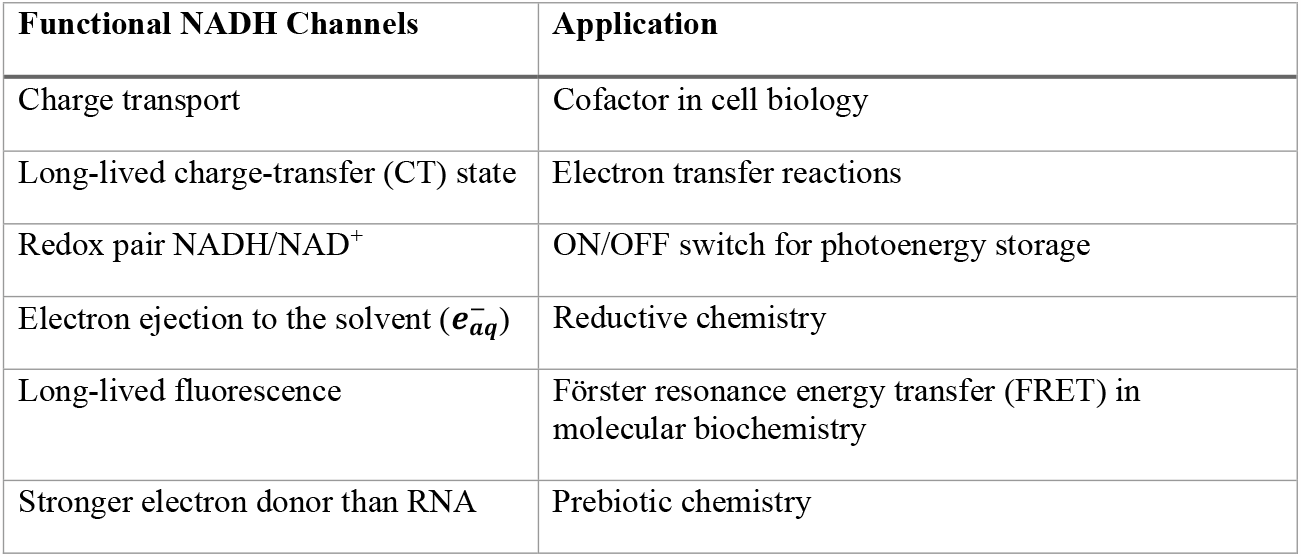
Overview of the diversity of molecular functions of NADH.

| Functional NADH Channels | Application |
| --- | --- |
| Charge transport | Cofactor in cell biology |
| Long-lived charge-transfer (CT) state | Electron transfer reactions |
| Redox pair NADH/NAD<sup>+</sup> | ON/OFF switch for photoenergy storage |
| Electron ejection to the solvent ($e_{aq}^-$) | Reductive chemistry |
| Long-lived fluorescence | Förster resonance energy transfer (FRET) in molecular biochemistry |
| Stronger electron donor than RNA | Prebiotic chemistry |

## ASSOCIATED CONTENT

### Supporting Information

Detailed materials, theoretical methods and methods on sample preparation and transient absorption measurements, supplementary results for NAD^+^.

## AUTHOR INFORMATION

The authors declare no competing financial interests.

## ACKNOWLEDGMENT

This work was supported in part by the Simons Foundation (SCOL 290360FY18 to D.D.S.) and by the National Science Foundation (Grant No. 1933505 to D.D.S.). M.J.J. acknowledges the support of a computational grant from Wrocław Centre of Networking and Supercomputing (WCSS). The authors acknowledge the

