## Supplementary Information for "The Unexpected Functional Diversity of Photoexcited NAD"

### Materials and Methods

#### Samples

The β-Nicotinamide adenine dinucleotide reduced, disodium salt (NADH, ≥98%) was purchased from MP Biomedicals and $NAD^+$ (~100%) from Roche Diagnostics, Germany. For ultrafast pump-probe spectroscopy, NADH and $NAD^+$ were dissolved in 50 mM deuterated ($D_2O$) phosphate buffer at pD 6.9 and kept at concentrations of several millimolar. The absorbance of NADH was 0.9 OD in both 267 nm and 339 nm excitation experiments and 2.7 OD for $NAD^+$ (267 nm excitation). The time-resolved experiments were performed in a flow cell with $BaF_2$ windows (3 mm) and a sample thickness of 100 µm. All measurements were conducted at a room temperature of 24°C under ambient oxygen conditions.

#### Transient Measurements

The basic principles of ultrafast pump-probe spectroscopy are described in the literature.[1-3] A Ti:Sa based laser amplifier system (Solstice Ace, Spectra Physics), with an output pulse duration of ~100 fs, 1 kHz repetition rate, 800 nm wavelength was used as light source. The 267 nm and 339 nm excitation pulses were generated in a non-linear amplifier system (Topas Prime + NIRUVis, Light Conversion, Ltd) and stretched to ~1.7 ps by a UV fused silica block (Corning, length 25 cm). The excitation pulses were focused to a pump spot diameter of ~110 µm FWHM at the sample position. The excitation energy at the sample position was between 0.88 µJ and 0.98 µJ. Energy dependence measurements did not reveal any quadratic conduct indicative of 2-photon excitation processes. The probe pulses were generated in a non-linear amplifier system (Topas Prime + DFG2, Light Conversion, Ltd) and focused to probe spot diameter of ~200 µm FWHM at the sample position inside a transient absorption spectrometer (Helios FIRE, Ultrafast Systems LLC). Pump and probe pulses were then spatially overlapped in the sample under magic angle conditions. The transmitted probe pulses and a second infrared beamline as a reference were spectrally dispersed (iHR 320, Horiba) and detected on two 64-channel MCT arrays (MCT-13-2x64, Infrared Associates Inc.). All mid-IR parts of the setup were purged with dry air. The transient lifetimes were determined from global fitting analysis.[4-6]

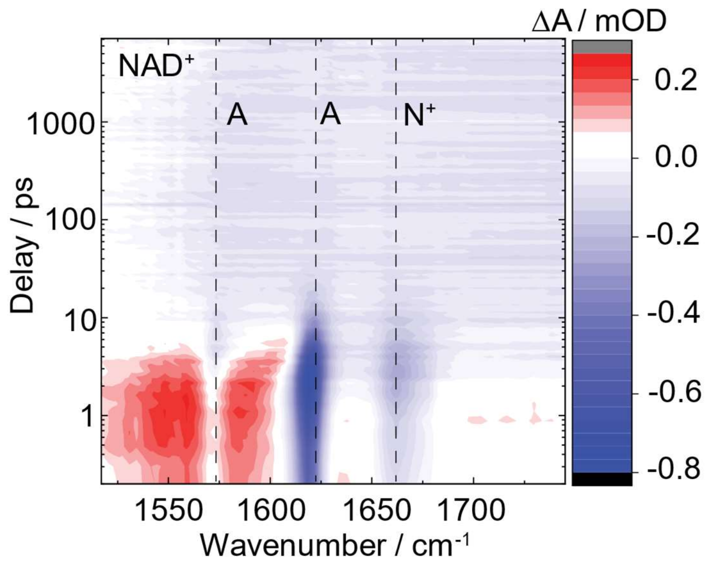


**Figure S1.** Transient 266 nm pump, mid-infrared probe absorbance difference spectra of $NAD^+$ in aqueous buffered (pD 6.9) solution, as a function of wavenumber (x-axis) and delay time (y-axis). Positive signals are shown in red and negative in blue.

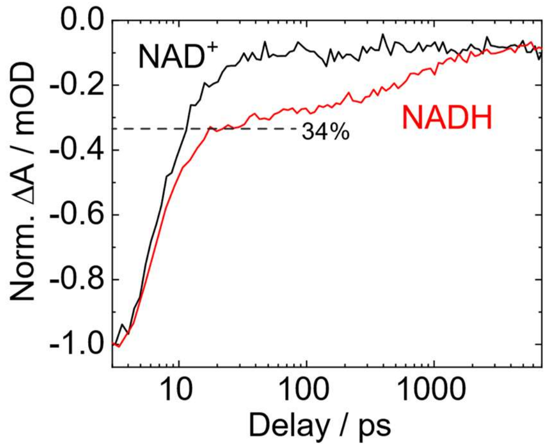


**Figure S2.** Time dependence of the mid-infrared absorbance differences of $NAD^+$ (black) and NADH (red) at the adenosine ground-state band around 1623 $cm^{-1}$ following excitation at 267 nm. For comparability, both plots were normalized at the local minimum around 3 ps. The rapid decay of both plots within the first 10 ps indicates vibrational cooling. In the case of NADH, a plateau which originates from the fluorescent and charge transfer decay channels is reached at a 34% level with a lifetime of several 100 ps.

### Theoretical section

### Computational methods

The starting ground-state structure of the folded NADH molecule was adapted from the classical molecular dynamics simulations conducted by Buckley et al.[7] To create the continuous solvation shell around the investigated molecule, 351 explicit water molecules were added to the mentioned NADH geometry. To surround the NADH molecule consistently using explicit water molecules, we employed the quantum cluster growth (QCG) procedure[8] implemented in the Conformer-Rotamer Ensemble Sampling Tool[9] (CREST). The QCG extension of the CREST was applied with the semi-empirical GFN2-xTB level of theory[10] accessible in the xtb program[11] package. To generate the water cluster employing the QCG tool, an automated computational interaction site screening (aISS) workflow[12] was utilized. Furthermore, the analytical linearized Poisson-Boltzmann (ALPB) model was used as an implicit solvent model, assuming water solvent. During the cluster growth, the initial NADH structure was completely frozen.

The obtained solvated NADH structure at the ALPB/GFN2-xTB level of theory (see Fig. S3), having 351 water molecules, was optimized using the ONIOM method[13] in the ORCA 5.0.3[14] program. The subtractive QM/QM2 coupling scheme was employed with the electrostatic embedding in the latter method. In the QM/QM2 calculations, the entire NADH molecule was described using the Kohn–Sham density functional theory (KS-DFT) and the range-separated hybrid ωB97X-D3 functional,[15] including improved dispersion correction[16] (DFT-D3), and together with the def2-TZVP basis set.[17] Whereas all explicit water molecules were treated by the semi-empirical GFN2-xTB method.[10] The equilibrium ground-state structure of NADH with 351 water molecules was optimized at the ONIOM(ωB97X-D3/def2-TZVP:GFN2-xTB) level of theory.

Vertical excitation energies of the extracted NADH structure from the optimized $S_0$ solvated system at the ONIOM(ωB97X-D3/def2-TZVP:GFN2-xTB) were computed using the algebraic diagrammatic construction to the second order method[18, 19] [ADC(2)] together with the COSMO implicit solvent model,[20] assuming the non-equilibrium variant and the def2-TZVP basis set[17] (COSMO-ADC(2)/def2-TZVP). Natural transition orbitals (NTOs) were generated to determine the molecular orbital character of energetically low-lying excited states. The charge transferred between the dihydronicotinamide and adenosine moiety was estimated employing the one-electron transition density matrix (1TDM) and the Löwdin style analysis. The NTOs and 1TDM were obtained using the TheoDore 3 package.[21]

The harmonic vibrational frequency analysis was performed for the equilibrium ground-state structures obtained using the ωB97X-D3 functional[15] with conductor-like polarizable continuum model[22] (CPCM) and the def2-TZVPPD basis set[23] (CPCM/ωB97X-D3/def2-TZVPPD). The CPCM/ωB97X-D3/def2-TZVPPD frequency calculations were performed for separate molecules such as adenosine, adenosine radical anion, dihydronicotinamide riboside and dihydronicotinamide riboside radical cation. For these molecules, the ribose configuration was adapted from corresponding fragments of the NADH structure. Since time-resolved mid-infrared (mid-IR) experiments were performed in deuterated water, all exchangeable hydrogen atoms of the mentioned molecules were replaced by deuterium in the frequency calculations. Based on the obtained theoretical results, the mid-IR spectra were prepared using Lorentz lineshape having a half-width of 15 $cm^{-1}$.

The crucial geometries on excited-state potential energy surfaces of the nicotinamide riboside molecule were found using the ADC(2) method together with the aug-cc-pVDZ[24] basis set (ADC(2)/aug-cc-pVDZ), assuming the equilibrium ground-state geometry found at the CPCM/ωB97X-D3/def2-TZVPPD level of theory. The minimum-energy crossing point (MECP) between the first excited and ground state was located using the sequential penalty constrained optimization function applied in the CIOpt package.[25] The MECP was optimized employing energies and analytical gradients obtained using the MP2/ADC(2)/aug-cc-pVDZ method. To conduct the MECP optimization, the CIOpt[25] and Turbomole 7.6[26] programs were interfaced. The potential energy profiles (PEPs) of the excited-state pathway were built using the image-dependent pair potential (IDPP) interpolation[27] between the $S_0$, $S_1$ and $S_1/S_0$ minimum-energy geometries. To estimate the energy difference between the ADC(2) $S_1$ minimum-energy structure and the corresponding ground state, the multi-state complete active space second-order perturbation theory[28, 29] (MS-CASPT2), based on the state-averaged complete active space self-consistent field (SA-CASSCF) method with the cc-pVDZ basis set[24] and the polarizable continuum model[22] in the equilibrium variant for the $S_1$ excited state. In the multireference calculations, the complete active space (CAS) was constructed of molecular orbitals (MOs) with natural orbital occupations in the range of 0.02–1.98,[30] and contained four occupied π MOs together with three virtual π* MOs (overall 8 electrons were correlated in 7 orbitals). The SA-CASSCF wave function was averaged over two lowest-lying electronic states. For the $S_1$ minimum-energy microhydrated structure, harmonic vibrational frequencies were computed at the ADC(2)/cc-pVTZ level of theory, assuming exchangeable hydrogen atoms were replaced by deuterium.

**Ground-state structure of solvated NADH**

In this paper, our aim was to investigate the UV-induced charge-transfer mechanism in the folded form of the NADH molecule, allowing for the formation of a long-lived charge-separated state. Therefore, we decided to focus on the photophysical properties of aqueous NADH in the stacked configuration, in which the adenine and dihydronicotinamide moieties are situated in a parallel way at a close distance (<4.0 Å). The initial NADH structure in the folded form was adapted from the classical molecular dynamics simulations,[7] and the selected geometry possessed the C3'-endo and C2'-endo ribose configuration for the dihydronicotinamide and adenosine moiety, respectively. To find a reliable solvation shell of the selected NADH geometry and a hydrogen-bond network between solvent and solute, we used the quantum cluster growth (QCG) tool[8] to add 351 water molecules systematically at the ALPB/GFN2-xTB level of theory. In turn, we performed the ONIOM(ωB97X-D3/def2-TZVP:GFN2-xTB) calculations in which the NADH structure was treated at the KS-DFT level of theory, whereas all explicit water molecules were described using the semi-empirical GFN2-xTB method. The optimized ground-state structure of the solvated NADH system is presented in Fig. S3. The extracted NADH geometry from the optimized solvated system is shown in Fig. S4. The proposed ground-state geometry-optimization protocol allowed us to obtain the folded form of NADH in which the average distance between the adenine and dihydronicotinamide moieties is around 3.65 Å. Additionally, the initial ribose configurations for both fragments of NADH are kept. The obtained ground-state structure of NADH and its geometrical features agree very well with previous computational studies on the conformational space of NADH.[7] It is worth adding that 20-55% of NADH molecules in aqueous solution exhibit the stacked orientation of adenine and dihydronicotinamide,[31] and the remaining molecules are in the open configuration without intramolecular interactions between the fragments.

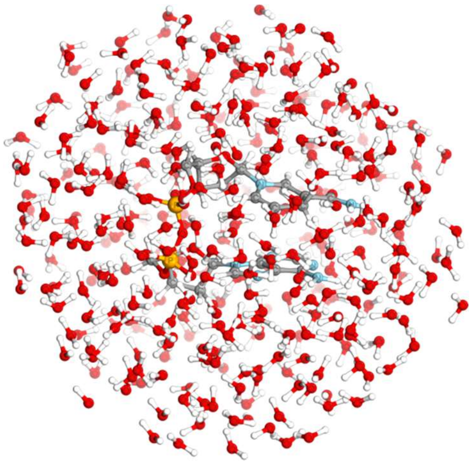

**Figure S3.** The optimized ground-state structure of the NADH molecule and 351 explicit water molecules found at the ONIOM(ωB97X-D3/def2-TZVP:GFN2-xTB) level of theory. All solvent molecules were treated by the semi-empirical GFN2-xTB method, and the entire NADH molecule was described at the KS-DFT level of theory.

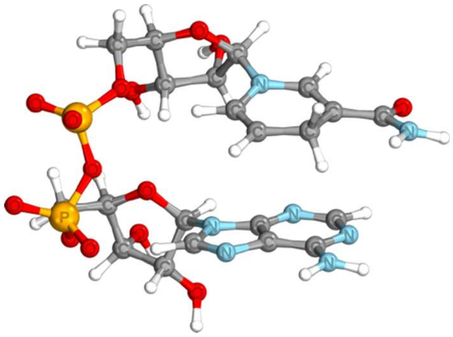

**Figure S4.** The extracted NADH geometry from the solvated NADH system optimized using the ONIOM(ωB97X-D3/def2-TZVP:GFN2-xTB) method.

### Photophysical properties of NADH

Although we consistently obtained the solvation shell alongside the hydrogen-bond network, the number of the added explicit water molecules (351) is insufficient to adequately describe the photophysical properties of the negatively charged NADH molecule in aqueous solution. Therefore, we took the NADH structure from the optimized solved system (Fig. S3) and used it as a target geometry for the following photophysical calculations. To account for solvent effects on the investigated molecule, we employed the COSMO implicit solvent model. Thus, vertical excitation energies of the NADH structure were obtained at the COSMO-ADC(2)/def2-TZVP level of theory, and the results are shown in Tab. 1.

Our COSMO-ADC(2) calculations predicted two optically bright states ($S_1$ and $S_4$) of the NADH molecule at 3.66 and 5.10 eV (see Tab. 1). The molecular orbital character analysis employing natural transition orbitals (see Fig. S5) revealed that both bright states have locally excited (LE) state character and are situated separately on the dihydronicotinamide (NH) and adenine (A) moiety. The $S_1$ and $S_4$ excited states are associated with the $^1\pi_{NH}\pi^*_{NH}$ and $^1\pi_A\pi^*_A$ electron transitions (Fig. S5), respectively. In addition, the $S_1(\pi_{NH}\pi^*_{NH})$ and $S_4(\pi_A\pi^*_A)$ states are characterized by a wavelength of 339 and 243 nm, respectively, and both wavelengths match very well with absorption maxima of the dihydronicotinamide (340 nm) and adenine (260 nm) fragments of NADH (Fig. 1). Since our theoretical calculations have correctly predicted the properties of the optically bright excited states of NADH compared to the experimental results (Fig. 1), the chosen computational protocol for studying the photophysical properties of NADH is credible. The $S_2$ excited state (4.81 eV), having the optically dark character, is marked by the $^1n_{NH}\pi^*_{NH}$ transition (Tab. S1) in which the $n_{NH}$ orbital is located on the carbonyl oxygen atom of the amide (Fig. S5). Surprisingly, among the energetically low-lying states, there is a charge-transfer (CT) $S_3$ excited state enabling the charge transfer of 0.74 $e^-$ from the dihydronicotinamide to adenine moiety, corresponding to the $^1\pi_{NH}\pi^*_A$ transition (Fig. S5). Interestingly, the $S_3(\pi_{NH}\pi^*_A)$ excited state has a non-negligible oscillator strength (0.089), suggesting that the CT state might be directly accessible by the absorption of UV photons. Consequently, the population of the CT $^1\pi_{NH}\pi^*_A$ excited state can form a charge-separated state in the form of the dihydronicotinamide radical cation and adenine radical anion.

**Table S1.** Vertical excitation energies (in eV) of the NADH molecule were obtained using the COSMO-ADC(2)/def2-TZVP level of theory, assuming the extracted NADH structure from the solvated NADH system found at the ONIOM(ωB97X-D3/def2-TZVP:GFN2-xTB) level of theory.

| State/Transition | | $E_{exc}$ [eV] | $f_{osc}$ | λ [nm] |
|---|---|---|---|---|
| $S_1$ | $^1\pi_{NH}\pi^*_{NH}$ (LE) | 3.66 | 0.1400 | 338.8 |
| $S_2$ | $^1n_{NH}\pi^*_{NH}$ (LE) | 4.71 | 0.0031 | 263.2 |
| $S_3$ | $^1\pi_{NH}\pi^*_A$ (CT) | 4.77 | 0.0893 | 259.9 |
| $S_4$ | $^1\pi_A\pi^*_A$ (LE) | 5.10 | 0.1670 | 243.1 |

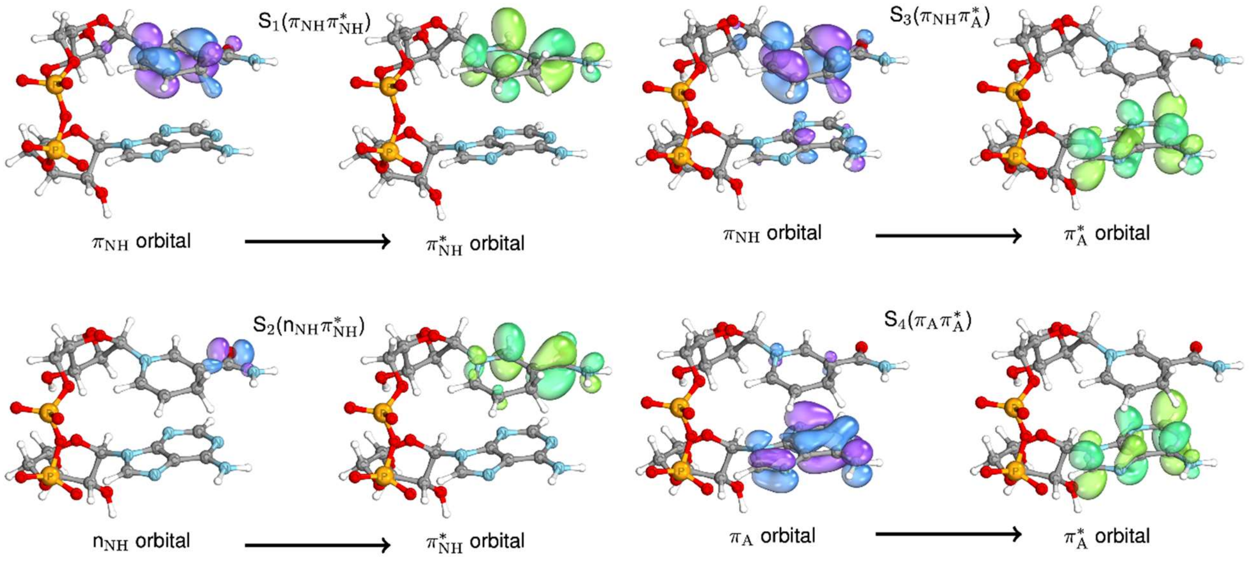


**Figure S5.** The natural transition orbitals created for the four lowest-lying excited states obtained at the COSMO-ADC(2)/def2-TZVP level of theory, assuming the extracted NADH geometry from the solvated NADH system optimized using the ONIOM(ωB97X-D3/def2-TZVP:GFN2-xTB) method.

**Photoinduced fluorescence mechanism of dihydronicotinamide riboside**

**Table S2.** Vertical excitation energies (in eV) of the dihydronicotinamide riboside structure were computed at the ADC(2)/aug-cc-pVDZ level of theory, assuming the equilibrium ground-state structure found at the ωB97X-D3/def2-TZVPPD/CPCM(Water) level of theory.

| State/Transition | | $E_{exc}$ [eV] | $f_{osc}$ | λ [nm] |
|---|---|---|---|---|
| $S_1$ | $^1\pi\pi^*$ | 3.82 | 0.1306 | 324.6 |
| $S_2$ | $^1\pi\sigma^*$ | 4.27 | 0.0026 | 290.4 |
| $S_3$ | $^1n_O\pi^*$ | 4.53 | 0.0003 | 273.7 |

To elucidate the photophysical and fluorescence properties of the dihydronicotinamide riboside structure constituting NADH, we performed excited-state calculations using the ADC(2)/aug-cc-pVDZ method and assuming the equilibrium ground-state structure (see Fig. S7) found at the ωB97X-D3/def2-TZVPPD/CPCM(Water) level of theory.

In Tab. S2, the three lowest-lying excited states of gas-phase dihydronicotinamide riboside are shown. In the first excited state (3.82 eV), electron transfer occurs from the occupied π to unoccupied $\pi^*$ orbital ($^1\pi\pi^*$), and both orbitals are localized on the six-membered ring of the dihydronicotinamide moiety (see Fig. S6). The $S_1(\pi\pi^*)$ state has the optically bright character and is responsible for a broad absorption band of NADH in the range of 300-360 nm (Fig. 1). Employing the aug-cc-pVDZ basis set, having a set of diffuse functions has enabled the identification of the repulsive $S_2$ excited state (Tab. S2) in which the electron is promoted from the occupied π orbital to the virtual repulsive $\sigma^*$ orbital (Fig. S6) delocalized around both hydrogen atoms at the C4 atom. It is worth adding that low-lying repulsive $^1\pi\sigma^*$ excited states can lead to the hydrogen atom abstraction process from chromophore mediated by solvent water molecules.[32]

The third excited state (Tab. S2) is associated with the $^1n_O\pi^*$ transition (Fig. S6) in which the lone electron pair ($n_O$) orbital is localized on the carbonyl oxygen atom of the amide and the unoccupied $\pi^*$ orbital is situated on the dihydronicotinamide moiety. Both described $S_1(\pi\pi^*)$ and $S_3(n_O\pi^*)$ excited states (Tab. S2) of dihydronicotinamide riboside are consistent in terms of the molecular orbital character and excitation energy with the previously discussed photophysical properties of the dihydronicotinamide moiety of NADH (Tab. S1), but there are a few minor discrepancies. In particular, the vertical excitation energy (3.82 eV) of the bright state is blue-shifted by 0.16 eV, and the difference arises from applying a double-ζ basis set (aug-cc-pVDZ) with comparison to the triple-ζ basis set (def2-TZVP) calculations (Tab. S1). Furthermore, the $^1n_O\pi^*$ excited state (4.53 eV) is red-shifted by 0.18 eV, and the results (in Tab. S2) were obtained without the presence of any implicit solvent model, whose inclusion destabilizes the excitation energy of the dark state.

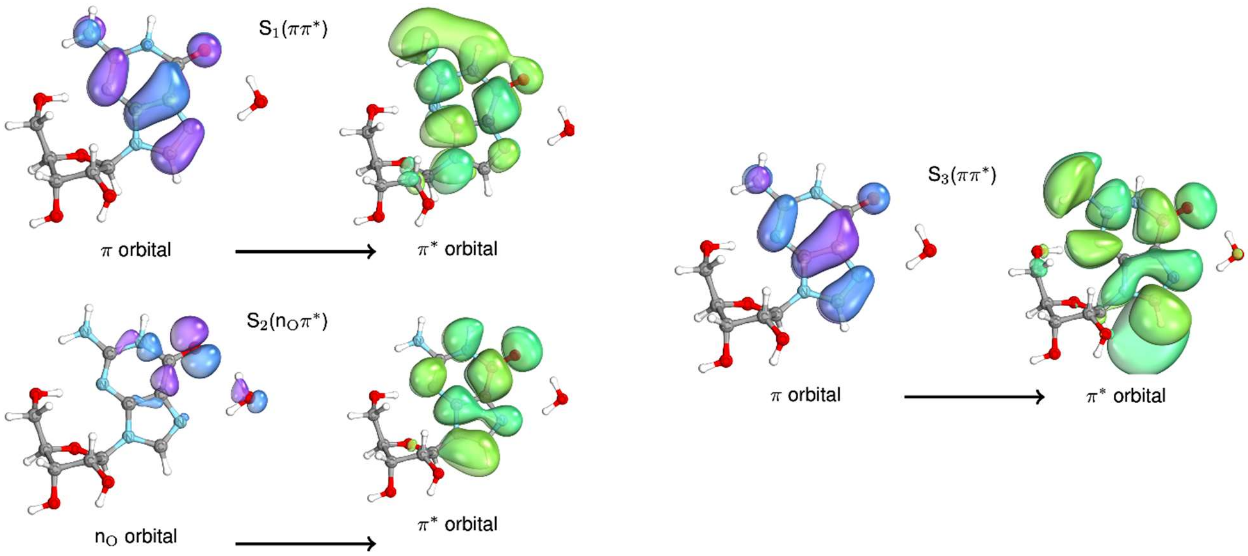


**Figure S6.** The natural transition orbitals were obtained for the three lowest-lying excited states computed at the ADC(2)/aug-cc-pVDZ level of theory, assuming the equilibrium ground-state dihydronicotinamide riboside structure found using the ωB97X-D3/def2-TZVPPD/CPCM(Water) method.

To study the photochemical pathways of UV-induced dihydronicotinamide riboside, we performed an exploration of the excited-state potential energy surfaces allowing for locating the $S_1(\pi\pi^*)$ minimum-energy structure and the $S_1(\pi\pi^*)/S_0$ minimum-energy crossing point using the MP2/ADC(2)/aug-cc-pVDZ method. Having the optimized $S_0$, $S_1$ and $S_1/S_0$ structures, we obtained the potential energy profiles (Fig. S7) from the Franck-Condon region through the $S_1$ minimum to the $S_1/S_0$ surface crossing.

Upon UV excitation, the system can directly populate the $S_1(\pi\pi^*)$ excited state (3.82 eV, lime line in Fig. S7). On its hypersurface, there is a barrierless pathway towards the $S_1(\pi\pi^*)$ minimum-energy structure (3.12 eV) that is characterized by the ring puckering occurring at the C2 atom (inset in Fig. S7), and the amide group is slightly out-of-plane of the six-membered ring. Subsequently, the UV-excited molecule can reach the sloped $S_1(\pi\pi^*)$ minimum-energy crossing point (inset in Fig. S7), marked by more pronounced C2 ring-puckering than in the $S_1$ minimum. Following the excited-state path, a sizable energy barrier of 0.68 eV between the $S_1$ minimum and

$S_1/S_0$ surface crossing appears. Thus, our results have shown that the UV-excited system must overcome the significant energy barrier to reach the $S_1/S_0$ crossing point, enabling radiationless deactivation to the corresponding electronic ground state. Hence, owing to the substantial energy difference between the presented $S_1$ and $S_1/S_0$ geometry, the system can stay in the $S_1$ minimum region for tens or hundreds of picoseconds. Therefore, we expect that UV-excited dihydronicotinamide riboside can undergo a radiative photorelaxation pathway through fluorescence process. To estimate the wavelength of photons emitted by the system upon UV excitation, we conducted the multireference SA-2-CASSCF(8,7)/MS-CASPT2/cc-pVDZ calculations together with the PCM solvation model in the equilibrium limit of the $S_1$ state, assuming the $S_1$ minimum-energy structure obtained at the ADC(2)/aug-cc-pVDZ level of theory. Our calculations concerning the energy difference between the $S_1$ minimum and corresponding $S_0$ state have revealed that the UV-excited system can emit photons having a wavelength of 475 nm (Fig. S7), and it very well agrees with previous experimental measurements of fluorescence spectra,[33] having the emission maximum of around 466 nm for aqueous NADH and dihydronicotinamide riboside.

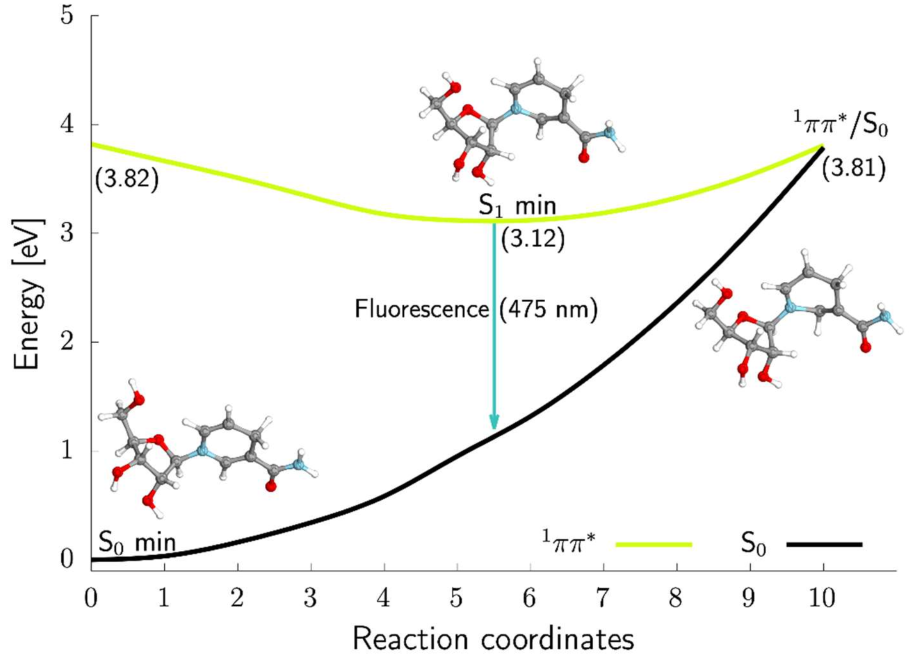


**Figure S7.** The potential energy profiles show the photochemical pathway of UV-excited dihydronicotinamide riboside from the optimized $S_0$ structure through the $S_1(\pi\pi^*)$ minimum and to the $S_1(\pi\pi^*)/S_0$ minimum-energy surface crossing (MECP), all these crucial geometries are added as insets. The energy barrier between the $S_1$ minimum and $S_1/S_0$ MECP amounts to 0.69 eV. The radiationless deactivation mechanism can occur when the system undergoes a C2 ring-puckering and out-of-plane displacement of the amide group. All excited and ground-state energies were obtained using the ADC(2) and MP2 methods, respectively, assuming the aug-cc-pVDZ basis set. The emission wavelength (475 nm) from the $S_1$ minimum was estimated based on the PCM(Water)/SA-2-CASSCF(8,7)/MS-CASPT2/cc-pVDZ calculations, and the implicit solvent model was in the equilibrium limit for the $S_1$ excited state.

### Harmonic vibrational frequency analysis of NADH

In order to provide a mechanistic explanation for the recorded time-resolved (TR) UV-pump mid-infrared (mid-IR) spectra obtained for NADH in deuterated water (Fig. 3), we conducted a vibrational frequency analysis for the equilibrium ground-state structures of C2’-endo adenosine and its radical anion as well as C3’-endo dihydronicotinamide riboside and its radical cation obtained at the ωB97X-D3/def2-TZVPPD/CPCM(Water) level of theory (Fig. S8). According to our discussed experimental-theoretical results in the article, upon 267 nm excitation of NADH, we expect the population of the charge-transfer excited state (Table 1) that results subsequently in the electron transfer from the dihydronicotinamide to the adenine moiety. Hence, dihydronicotinamide riboside radical cation and adenosine radical anion are formed. The performed ground-state geometry optimization and a spin population analysis for both radicals show that an unpaired electron is delocalized over the six-membered ring of the dihydronicotinamide fragment and, in the case of adenosine, is shared between the C8 and C6 atoms.

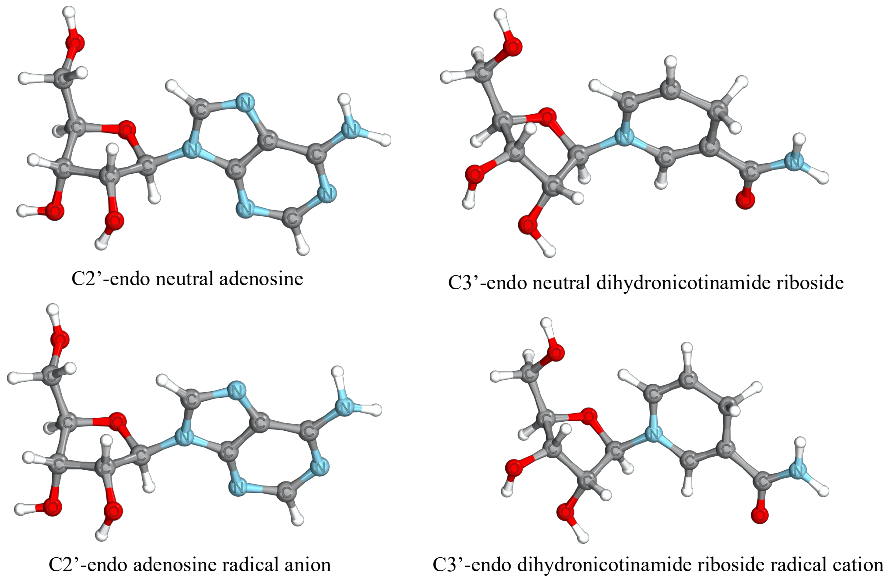


**Fig. S8** The presented equilibrium ground-state structures of neutral adenosine and its radical anion and neutral dihydronicotinamide riboside and its radical cation were obtained using the ωB97X-D3/def2-TZVPPD/CPCM(Water) method.

Table S3 presents our recorded TR-IR signals in the 1510-1750 $cm^{-1}$ region obtained during UV-pump mid-IR measurements of NADH in deuterated water. The experimental signals were assigned to specific vibrational modes based on the vibrational frequency analysis for the structures presented in Fig. S8. According to the experiment, there were recorded characteristic ground-state bleaching (GSB) bands at 1553 $cm^{-1}$, 1685 $cm^{-1}$, and 1624 $cm^{-1}$, and all of them were assigned to the dihydronicotinamide moiety. The acquired GSB signals at 1575 $cm^{-1}$ and 1624 $cm^{-1}$ originated from the adenosine part. The ground-state bleaching bands and their corresponding characteristic vibrational modes for NADH are consistent with previous studies.[34] However, a significant energy difference exists for computed symmetric (1773 $cm^{-1}$) and antisymmetric (1685

$cm^{-1}$) C2-C3 and C5-C6 bonds stretching and their experimental counterparts at 1685 $cm^{-1}$ and 1553 $cm^{-1}$, respectively. We may only speculate that the pronounced energy difference between the experimental and theoretical results could come from specific vibrational modes obtained for separate NADH chemical fragments (Fig. S8), which cannot interact with each other as in the NADH structure. One of the possible solutions for the problem would be performing ground-state dynamics of NADH molecule to obtain the IR spectrum and clarify whether any intramolecular interactions between the dihydronicotinamide and adenine ring could lead to more accurate frequencies for the C2-C3 and C5-C6 bonds stretching.

Alongside the GSB bands, there were also obtained long-lived positive (absorption) signals at 1510 $cm^{-1}$ and 1610 $cm^{-1}$ assigned to adenosine radical anion, and 1654 $cm^{-1}$ and 1726 $cm^{-1}$ corresponding to dihydronicotinamide riboside radical cation. The obtained experimental-theoretical results are very consistent (Tab S3). Furthermore, these TR-IR absorption bands are signatures of the long-lived charge-separated state that occurs after the population of the charge-transfer state (Table S1). It is worth adding that the absorption signals at 1553 $cm^{-1}$ and 1575 $cm^{-1}$ can cover the theoretically predicted positive band at 1562 $cm^{-1}$ corresponding to dihydronicotinamide riboside radical cation, and owing to that, the latter band is not visible in the experimental spectra (Fig. 3B)

**Table S3.** Experimental vibration bands in the 1510-1750 $cm^{-1}$ range obtained from the time-resolved (TR) UV-pump mid-IR measurements of NADH in deuterated water. Hamonic vibrational frequencies in the mid-IR region were computed at the ωB97X-D3/def2-TZVPPD/CPCM(Water) level of theory. In parentheses, specific vibrational modes are given.

| Adenosine (bleaching bands) [$cm^{-1}$] | | Adenosine radical anion (absorption bands) [$cm^{-1}$] | | Dihydronicotinamide riboside (bleaching bands) [$cm^{-1}$] | | Dihydronicotinamide riboside radical cation (absorption bands) [$cm^{-1}$] | |
|---|---|---|---|---|---|---|---|
| Theory | TR-IR | Theory | TR-IR | Theory | TR-IR | Theory | TR-IR |
| 1539 (Symmetric C6-$NH_2$ and N3-C2 stretching) | 1575 | 1520 (Antisymmetric N9-C4 and C4-N3 stretching) | 1510 | 1704 (Antisymmetric C2-C3 and C5-C6 stretching) | 1553 | 1562 (Antisymmetric C2-C3 and C5-C6 with N1 stretching) | - |
| 1662 (six-membered ring stretching with C6-$NH_2$ bond stretching) | 1624 | 1600 (Symmetric C4-C5 and N1-C2 stretching) | 1610 | 1625 (C-O stretching) | 1624 | 1681 (Symmetric C-O, C2-C3 and C5-C6 stretching) | 1654 |
| - | - | - | - | 1773 (Symmetric C2-C3 and C5-C6 stretching) | 1685 | 1726 (Antisymmetric C-O stretching with symmetric C2-C3 and C5-C6 stretching) | 1726 |

Direct excitation of the dihydronicotinamide (NH) moiety of NADH induces fluorescence, therefore, we expect that upon selective excitation, the NH fragment should be found in the lowest-lying excited state, according to the Kasha's rule,[35] and the experimentally tracked photoinduced dynamics of NADH should mainly occur in the $S_1$ excited state of the molecular fragment. To explain the origin of a long-lived signal below 1525 $cm^{-1}$ acquired during the time-resolved UV-pump (at 339 nm) IR-mid measurements at (Fig. 3A), we performed a vibrational frequency analysis for the optimized $S_1(\pi\pi^*)$ structure of the methylated dihydronicotinamide molecule with five quantum-chemical water molecules, which saturate all hydrogen bonds between the chromophore and solvent. All exchangeable hydrogen atoms were replaced by deuterium in the mentioned analysis. The $S_1$ minimum-energy structure was found at the ADC(2)/cc-pVTZ level of theory, and its structural features are in line with the similar $S_1$ minimum yielded for the dihydronicotinamide riboside structure (Fig. S7). To reduce the cost of calculations, we replaced the sugar moiety with the methyl group, and we added five quantum-chemical molecules to include explicit solvent interactions with the amide group in the excited-state calculations. The obtained harmonic vibrational frequencies for microhydrated methylated dihydronicotinamide in the $S_1$ minimum show that in the 1500-1800 $cm^{-1}$ region, there is only an intense absorption band at 1506 $cm^{-1}$. The main contribution to the band is associated with asymmetric stretching of the amide group. Importantly, these results correlate perfectly with the experimental TR-IR spectrum (Fig. 3A) in which there is the recorded positive signal below 1525 $cm^{-1}$, and there are two more absorption bands at 1654 $cm^{-1}$ and 1726 $cm^{-1}$, but they were previously assigned to the dihydronicotinamide radical cation structure.

Since the population of the $S_1$ minimum of the dihydronicotinamide moiety of NADH can last up to hundreds of picoseconds due to the lack of an efficient barrierless non-radiative deactivation pathway (Fig. S7), the UV-excited chromophore can emit photons by means of fluorescence process and deactivate itself to the electronic ground state. Therefore, we postulate that the recorded long-lived signal below 1525 $cm^{-1}$ can be assigned to the simulated absorption band at 1506 $cm^{-1}$ in the $S_1$ minimum. Hence, the experimentally observed signal should be a signature of the $S_1(\pi\pi^*)$ state that experiences fluorescence (Fig. S7) from the dihydronicotinamide moiety of NADH.

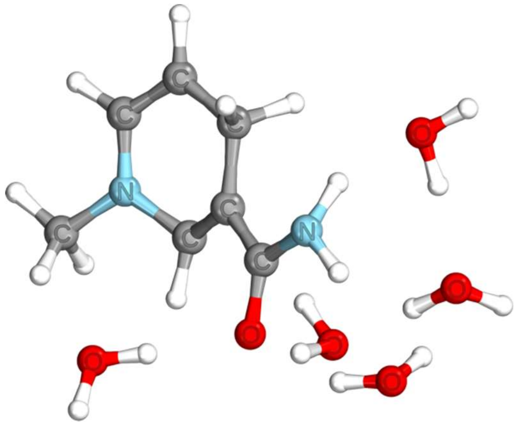


**Fig. S9** The shown $S_1(\pi\pi^*)$ minimum-energy structure of methylated dihydronicotinamide with five quantum-mechanical water molecules was optimized at the ADC(2)/cc-pVTZ level of theory.
